# High resolution profile of body wide pathological changes induced by abnormal elastin metabolism in Loxl1 knockout mice

**DOI:** 10.1101/353169

**Authors:** Bingbing Wu, Yu Li, Chengrui An, Deming Jiang, Lin Gong, Yanshan Liu, Yixiao Liu, Jun Li, HongWei Ouyang, XiaoHui Zou

## Abstract

Abnormal ECM caused serious body wide diseases and elastin is one of the important ECM components. But its systemic function still has not yet been thoroughly illustrated due to limitations related to novel research technologies. To uncover the functions of elastin, a new method for body-wide organ transcriptome profiling, combined with single-cell mass cytometry of the blood, was developed. A body-wide organ transcriptomic (BOT) map was created by performing RNA-seq of 17 organs from both *Loxl1* knockout (KO) and wide type (WT) mice. The BOT results showed a systematic up-regulation of genes related to immune response and proliferation process in multiple tissues of the KO mice; histological and immune staining also confirmed the hyperplasia and infiltration of local immune cells in the vagina, small intestine, and liver tissues of KO mice. Furthermore, using 32 markers, CYTOF mass cytometry analysis of the immune cell subpopulations from the peripheral blood revealed apparent systemic immune changes in the KO mice; data showed an activated NK cells and T cells with a higher expression of CD44 and CD38, and a suppressed B cells, macrophages and neutrophils with lower expressions of CD62L, CD44 and IL6. More interestingly, these findings also correlated well with the data obtained from cancer patient databases; tumor patients had higher mutation frequency of Loxl1, and the Loxl1-mutant tumor patients had up-regulated immune process, cell proliferation and decreased survival rate. Thus, this research provided a powerful strategy to screen body-wide organ functions of a particular gene; the findings also illustrated the important biological roles of elastin on multiple organ cells and systemic immunity. These strategy and discoveries are both of important value for the understanding of ECM biology and multi-organ cancer pathology.

## Introduction

The extracellular matrix (ECM) provides an essential microenvironment ingredients to support the integrity of tissues and maintain tissue homeostasis^1^. Various components of ECM such as hyaluronic acid, tenascin-C, fibronectin, and agrin could differentially regulate cells’ fate and functions (DNA synthesis, proliferation, migration, and differentiation) during regeneration and homeostasis ^2–4^ Additionally, the ECM physical properties such as mechanical strain, substrate stiffness, shear stress and topography can also affect stem cell fate determination ^5^.

Disrupted ECM homeostasis or dysregulated ECM remodeling is correlated with pathological conditions and can worsen disease progression. ECM disruption (collagen, elastin, hyaluronic acid) can lead to alternations in junctional integrity, and influence vascular permeability and the release of inflammatory factors ^6^; thus, these processes are related to diverse body-wide pathological conditions and diseases including embryonic lethality ^7^, inflammatory response and fibrosis ^8^, aging^9^, and cancer^10^.

Elastin is one of the key ECM protein in connective tissues that provides elasticity and reversible deformability ^11^. The loss of elastic tissue (elastin) could lead to serious organ deformities such as cutis laxa ^12^, Supravalvar aortic stenosis, Williams syndrome^13^, and bullae^14^, highlighting the importance of elastin on organ morphology maintenance; however, the biological functions of elastin in body-wide organ cell activities have not yet been well-characterized.

The current strategy involving the use of gene knockout mice to investigate the biological functions of a particular gene is mainly focused on the visible phenotype of limited organs. By using Loxl1 (an enzyme responsible for elastin synthesis and cross-linking)^15,16^ knockout mice (KO mice), previous studies illustrated that Loxl1 deficiency led to abnormal elastin regulation, and subsequently resulting in pelvic organ prolapse (POP), flabby skin, and bullae ^14^ However, because of the body-wide expression of elastin ECM, its biological effects and clinical relevance in other organs were always ignored.

Due to significant advances in high-throughput sequencing technologies, biological functions of a single gene could be deciphered at whole genome and single cell resolution ^17^ Thus, in this study, we developed an efficient body-wide organ transcriptomic profiling method combined with single-cell mass cytometry to evaluate biological functions of abnormal elastin regulation through single-cell mass cytometry and RNA-seq of 17 organs from both abnormal (Loxl1 knockout (KO)) and normal (wide type (WT)) elastin mice.

## Results

### 1. Abnormal elastin-associated phenotype and other systemic changes in KO mice

Because Loxl1 is the key enzyme for the synthesis and assembly of elastin, the KO mice displayed phenotypes associated with abnormal elastic fibers, including obvious POP (figure1A) and loose skin (figure1B); POP is characterized by the enlarged perineal body and a bulge of rectum and vagina. In human, the molecular mechanism of POP is widely considered to be related to the theory of “imbalance of elastin” ^18^; thus, Loxl1 knockout mice are always used as the animal model for studies of POP. In order to observe changes of elastic fibers in KO mice, we performed weigarts’ staining specific for elastic fibers on vaginal tissues (figure1C) and skin (figure1D). In the KO mice, elastic fibers in the lamina propria of vagina and skin appeared to have a thin and short rod-shaped structure; additionally, we also observed that the deposition of elastic fibers near the basement membrane decreased in the KO mice model. On the contrary, elastic fibers were polarized and arranged neatly in the vagina tissues of WT mice, confirming the reduction of synthesis and failure of crosslinking process of elastic fibers in KO mice. Moreover, the length and volume of the spleen were significantly increased in the KO mice group when compared with the WT group (figure1E). The body weight of the KO mice was also measured where results revealed a significant weight decrease when compared to the WT group with increasing age (figure1F). These additional phenotypic changes suggested that the impact of Loxl1 deficiency on mice was not limited only to the reproductive tracts, but also on spleen and other parts of the body.

**Figure1.**
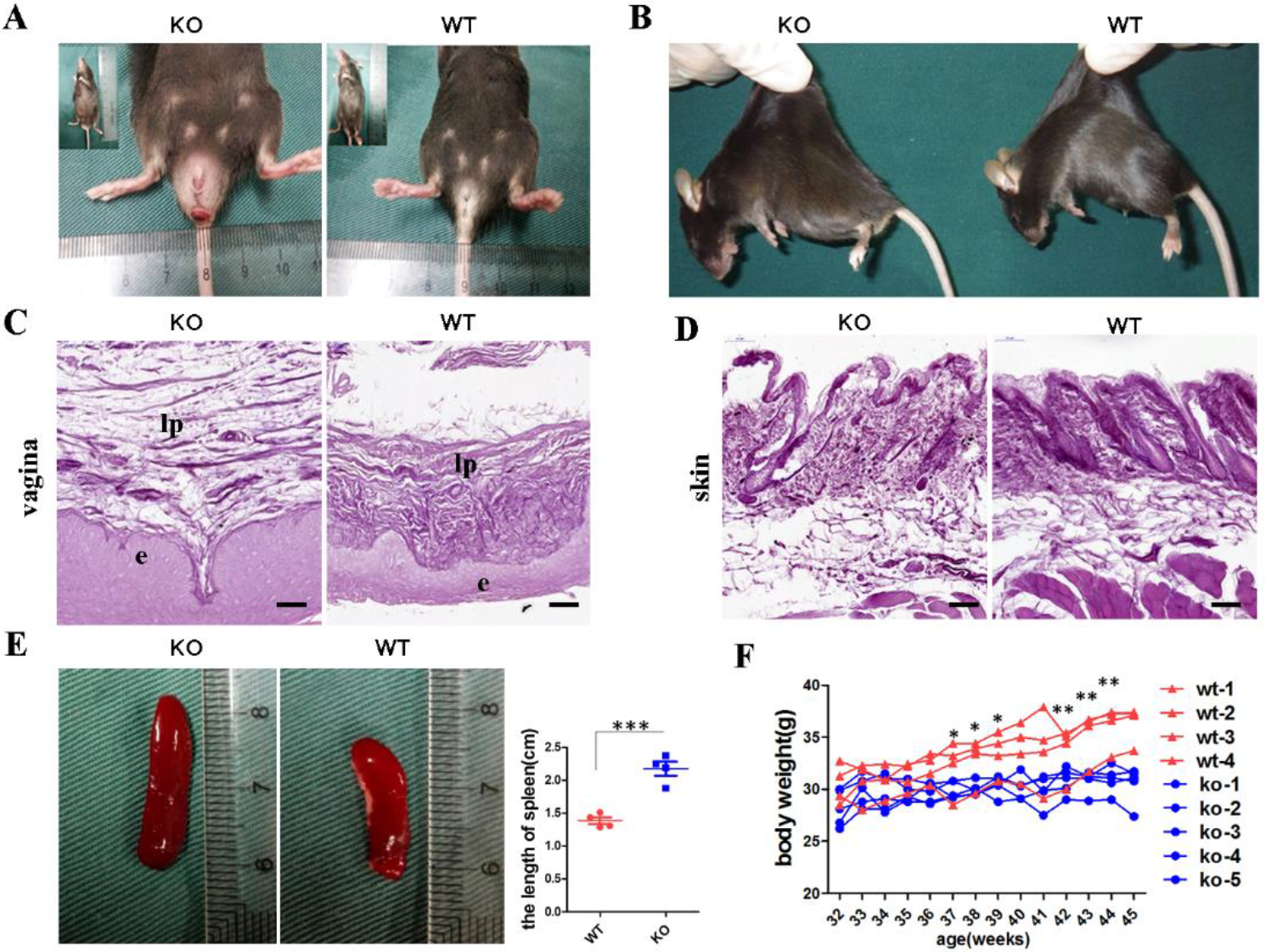
Abnormal elastin-associated phenotype and other systemic changes in KO mice. **A.** Overview of phenotype of pelvic floor in the Loxl1 knockout (left) and wild type mice (right). **B.** Overview of the loose skin in KO mice. **C, D.** Weigarts’ staining for elastic fiber in vagina (**C**) and skin (**D**), scale bar, 50 μm. **E.** Left: overview of the spleen. Right: quantitation of the length of spleen, the data represent mean ± s.d. of four samples. ***P<0.001. **F.** The body weight of the Loxl1 knockout and wild type mice, the line represent data of each mouse. **P<0.01, *P<0.05.

### 2. Up-regulated transcriptomic level of inflammation and proliferation in KO mice through body-wide organ RNA-seq

In order to assess the systemic effect of abnormal elastin on body-wide organs and tissues with the use of Loxl1 KO mice, we performed RNA-seq of 17 tissues (heart, liver, spleen, lung, kidney, skin, brain, aorta, abdominal fat, small intestine, bladder, rectum, vagina, skeleton muscle, bone, cartilage, and tail tendon) isolated from WT and KO mice with biological triplicate in each group (figure2A). We first employed a correlation analysis of gene expression across all different samples in the WT group; data showed that there were high correlation between the expression profiles of the same tissue types. The closely-related tissues such as rectum and small intestine, cartilage and bone, and skeleton muscle and skin can also be clustered together respectively according to the gene expression, indicating good repeatability in sequencing results (figure2B). The principle components analysis was also performed; figure 2C shows a tSNE of this data demonstrating clear separation in gene expression profiles between the KO and WT mice group, implying the differential characteristics of the two groups (figure2C).

To further compare the differentially expressed genes from the two groups in various tissues, we used the DESeq2 in R (http://www.R-project.org) for downstream analyses (figure2D). Data revealed that the spleen had the largest number of significantly differentially expressed (DE) genes (P value< 0.05), followed by vagina and skeleton muscle, thus suggesting that Loxl1 deficiency has the greatest effect on spleen. According to the generated heatmap of all DE genes in all tissues (figure2E), we found that most DE genes were unique to each tissue, possibly implying that Loxl1 deficiency has tissue-specific effects; gene ontology analysis (https://david.ncifcrf.gov/) of DE genes in different tissues also further verified their tissue-specific changes after Loxl1 knockout. For example, the up-regulated DE genes in the spleen were enriched with genes related to cell cycle and cell division after Loxl1 knockout, while the down-regulated DE genes were involved in immune system process (figure2F). In contrast, genes related to the biological processes of keratinization and keratinocyte differentiation were up-regulated, while sarcomere organization and response to estradiol were down-regulated in the vagina (supplementary figure1A). In skeletal muscles, the up-regulated GO terms included mitotic nuclear division and mitotic cell cycle, and the down-regulated GO terms comprised of immune system process and antigen processing and presentation (supplementary figure1B). Alterations of biological processes in other tissues were presented in supplementary figure2.

In addition to the tissue-specific changes in different tissues, we identified 38 common DE genes in more than 5 tissues (figure2G); GO enrichment of cell component analysis showed that the common up-regulated genes were mainly associated with extracellular exosomes (20/38) (e.g., COL1A1, COL6A3, COL1A2,H2-K1, ACTB, IFITM3, and *FBN1*), cytoplasmic components (23/38) (e.g., *ZFP36*, *TXNIP*, *EGR1*, *ACTB*, *CRIP1*, *IFITM3*, *MNDAL*, *SNCA*, and *NFKBIA*), nucleus components (17/38) (e.g., *ZFP36*, *TXNIP*, *EGR1*, *IFITM3*, *SNCA*, *MNDAL*, and *SUN2*), and basement membranes (4/38) (e.g., *ALB*, *FBN1*, *SPARC*, and *LOXL1*). Addition, the GO term of biological process analysis also revealed that these DE genes were enriched in “immune system process”, “response to bacterium”, and “cell proliferation” (figure2H); thus, these indicated that multiple tissues had common significant changes in the immune process and cell proliferation at the transcriptomic level.

**Figure2.**
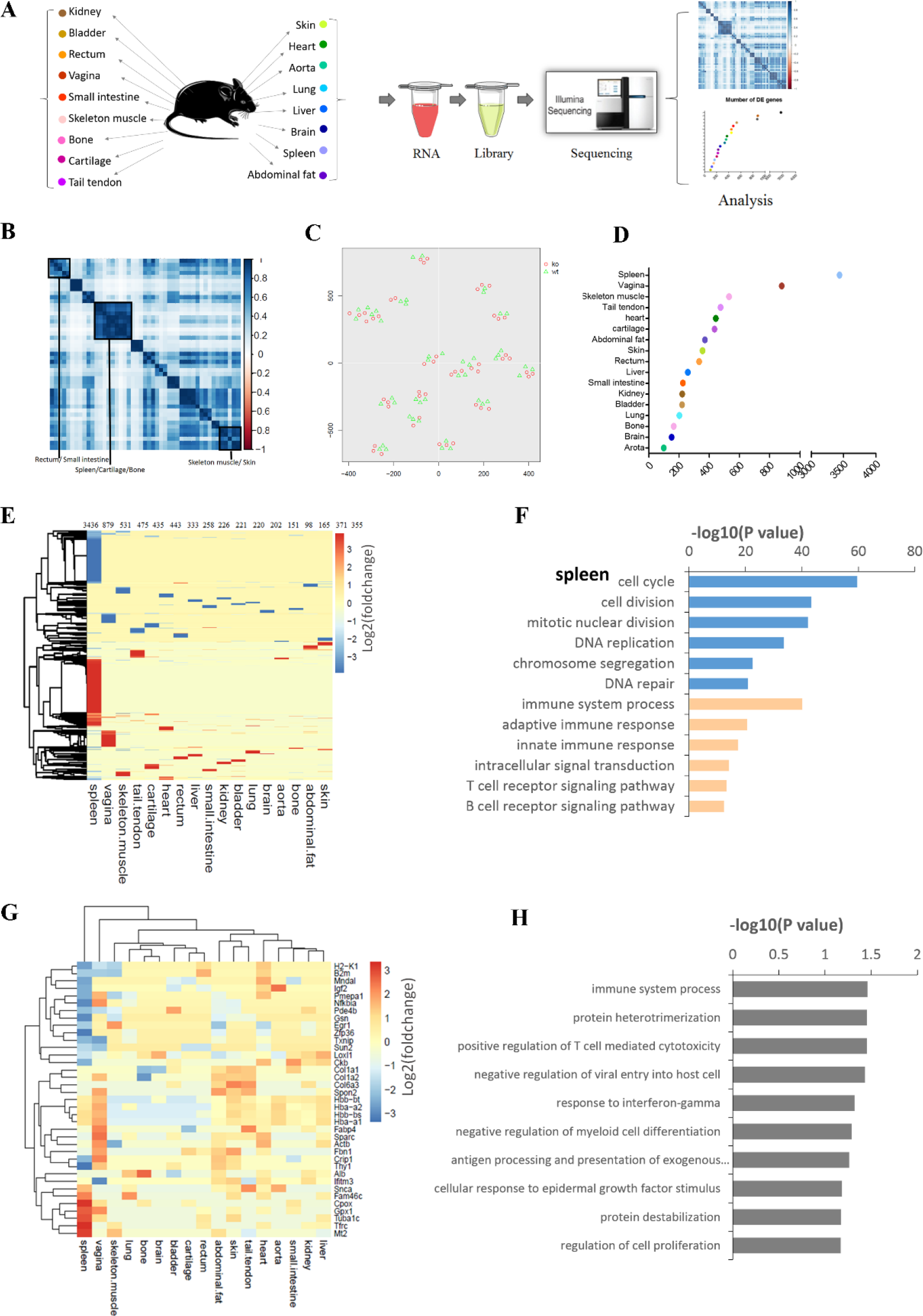
Up-regulated inflammation and proliferation in transcription level through body wide organ RNA-seq. **A.** scheme of the body wide organ RNA-seq, there were three animals in KO and WT groups and 17 organs were isolated in each animal. **B.** Heatmap of the pairwise correlation between all 17 tissues from WT group based on transcript expression levels of 46,814 genes. **C.** tSNE clustering analysis of the body wide organ mRNA profiles for KO and WT groups. **D.** Scatter plot of number of significantly differently expressed (DE) genes in 17 tissues. **E.** Heatmap of the DE genes in all tissues ordered by hierarchical clustering. Shown are log2(foldchange) values relative to compartment in WT group (P value< 0.05). **F.** Biological process gene ontology (GO) analysis of DE genes in spleen using DAVID software. The GO terms in blue color were up-regulated gene sets in KO group compared to WT group and the terms in orange color were down-regulated ones. The six most significant gene sets per condition were shown. **G.** Heatmap of the common DE genes (shared in ≥ 5 tissues) ordered by hierarchical clustering. Shown are log2(foldchange) values relative to compartment in WT group (P value< 0.05). **H.** Biological process GO analysis of common DE genes using DAVID software.

**Supplement figurel.**
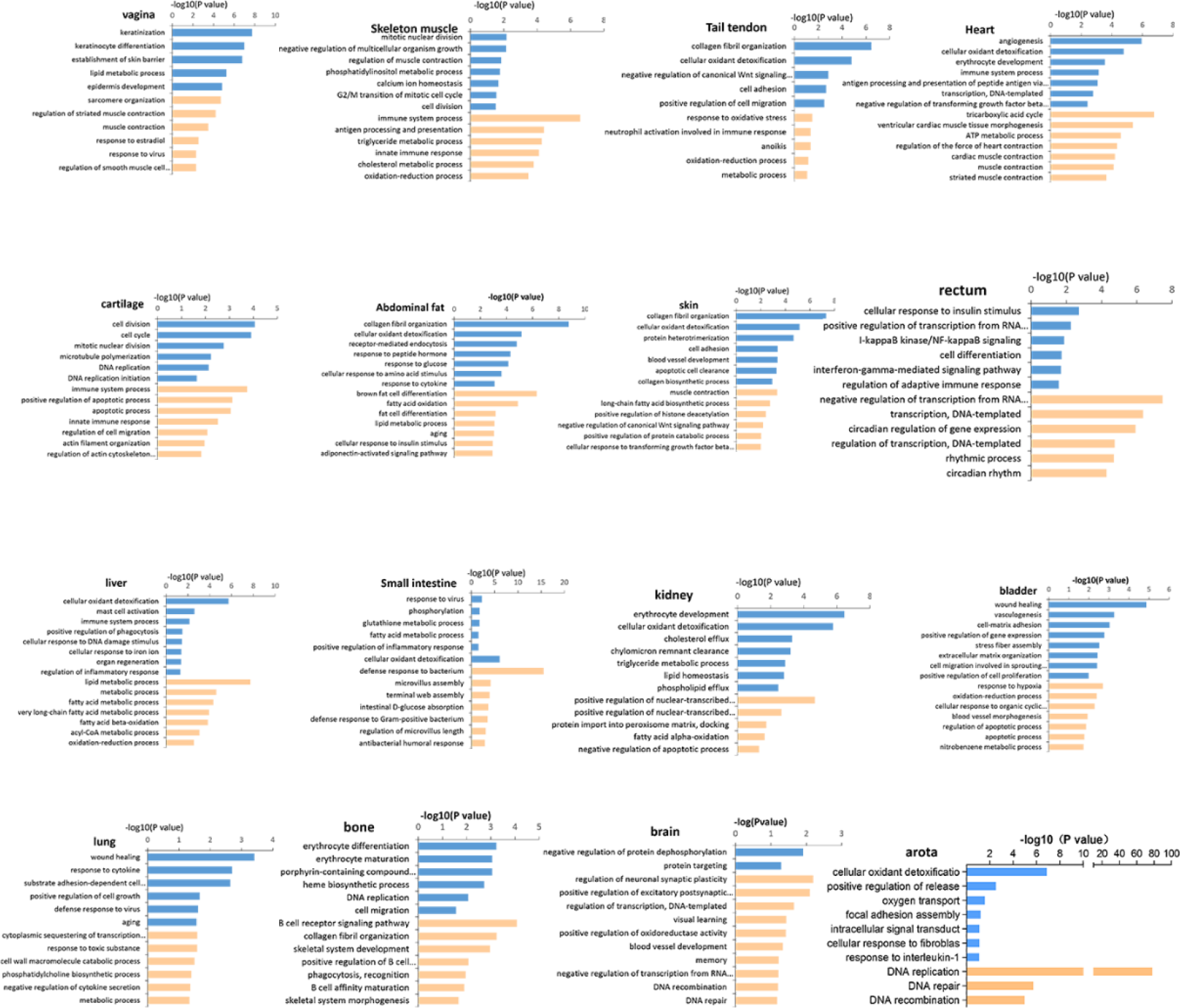
Biological process gene ontology (GO) analysis of DE genes in 16 tissues using DAVID software. The GO terms in blue color were up-regulated gene sets in KO group compared to WT group and the terms in orange color were down-regulated ones.

### 3. Tissues hyperplasia and local immune cells infiltration in KO mice

To verify that changes of cell proliferation occurred in the KO mice, histological staining was performed to further analyze all tissue samples. Haemotoxylin and Eosin(H&E) staining demonstrated that the vaginal epithelium appeared to be thicker, and the villi of the small intestine were longer when compared to the WT group (figure3A); on the contrary, other tissues including the skin, rectum, and liver showed no apparent changes when compared between the two groups.

Next, we performed immunofluorescence staining with a proliferation marker KI67. Results showed that the vaginal basal lamina, liver, and crypts of small intestine of the KO mice had increased expression of KI67 when compared with the WT mice (figure3B, C); these results were consistent to the previous transcriptomic data which showed an up-regulation of GO terms related to cell proliferation (figure3D) including keratinization, epidermal cell differentiation, and positive regulation of mesenchymal cell proliferation. Immunofluorescence staining of KI67 was also repeated in other remaining tissues; however, no differences were observed between the two groups.

Subsequently, immunofluorescent staining of anti-CD45, anti-CD19, and anti-F4/80 was also performed to detect local infiltration of immune cells. In the KO mice group, the number of CD45 positive cells, CD19 positive cells, and F4/80 positive cells increased in the vagina and small intestine (figure3E), indicating the local infiltration of immune cells (leukocyte, B cells, and macrophage). These results could match the transcriptomic data with an up-regulated GO terms which were enriched in immune process (figure3F). According to above results, it can be concluded that Loxl1 deficiency would cause hyperplasia and local immune cell infiltration in some tissues.

**Figure3.**
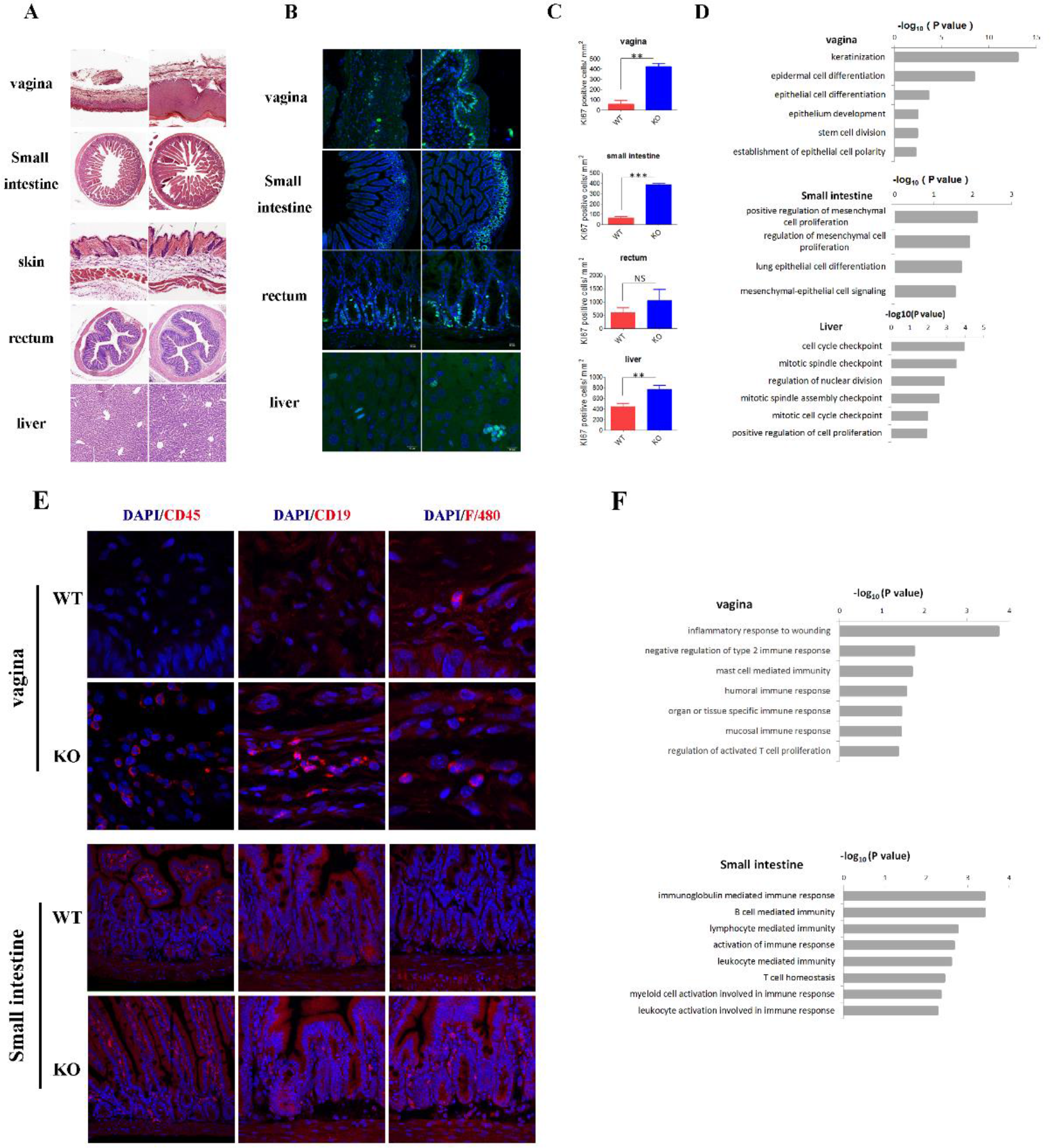
Tissues hyperplasia and local immune cells infiltration in KO mice. **A.** H&E staining of multiple tissues in WT and KO mice, including vagina, small intestine, skin, rectum and liver. **B.** Immunofluorescence staining of KI67 in tissues of vagina, small intestine, rectum and liver in WT and KO mice. **C.** Bar graphs of the numbers of KI67 positive cells per mm^2^ in tissues of intestine, skin, rectum and liver. The data represent mean ± s.d. of three samples. ***P<0.001, **P<0.01, N.S., no significance. **D**. Cell proliferation related gene ontology (GO) of up-regulated DE genes in tissues of vagina, small intestine and liver of KO mice. **E**. Immunofluorescence staining of immune cell makers in tissues of vagina, small intestine. **F.** Upregulated immune process related gene ontology (GO) in tissues of vagina and small intestine of KO mice.

### 4. Systematic immune changes in KO mice detected by CYTOF

Due to the up-regulation of immune-related GO terms detected in RNA-seq, we predicted that there would also be changes in the systemic immunity in KO mice. Therefore, to evaluate this point, we applied CYTOF mass cytometry to characterize the immune populations of the peripheral blood from WT and KO mice using 32 cell markers, in order to assess the impact of Loxl1 inefficiency on systemic immunity. The antibody panel contained major innate and adaptive immune cell subset markers, including cell surface markers, function makers, and cytokines. The CYTOF data was processed with Cytobank (www.cytobank.org) and SPADE, tools that were developed for visualization of complex cell composition and proportion in the peripheral blood (figure4A).

Well-recognized cell surface markers were utilized to identify specific cell subsets in the peripheral blood (figure4B). We first gated to obtain CD45+ cells in the peripheral blood (supplement figure2A); SPADE showed that the gated cells were all CD45 positive (supplement figure2B). We then identified major cell subpopulations within the CD45+ cell population (figure4B, C); among the CD3+ cell, we identified cells that highly expressed CD4, CD8 and gdTCR as CD4+ T cells (CD3+CD4+), CD8+T cells (CD3+CD8+), and gd T cells (CD3+gdTCR+), respectively. We also identified B cells (CD45+IgM+) and four subsets of highly-expressed CD11b cells, including neutrophils (CD45+CD11b+Ly6G+), monocytes (CD45+CD11b+Ly6C+), macrophages (CD45+CD11b+F4/80+), and NK cells (CD45+CD11b+CD49b+).

The proportion of each immune cell subset to the whole CD45+ cells of WT and KO group was then compared to assess the effect of *Loxl1* knockout on the immune system (figure4D). The WT and KO groups had the same proportion of CD3+ T cells (WT, 13.47%; KO, 13.51) and monocytes (WT, 6.65%; KO, 6.07%) while the ratio of CD4+ T cells (WT, 5.00%; KO, 3.87%) and B cells (WT, 19.03%; KO, 11%) showed a slight decrease in the KO group. Moreover, CD8 T cells (WT, 7.67%; KO, 9.05%), gd T cells (WT, 0.24%; KO, 0.39%), neutrophil (WT, 43.11%; KO, 50.76%), macrophage (WT, 4.75%; KO, 8.11%), and NK cells (WT, 8.65%; KO, 11.11%) had a relatively higher proportion in the KO compartment when compared to the peripheral blood of WT mice. These results illustrated that Loxl1 deficiency influenced the immune system in mice as evidenced by alterations in the proportion of immune cell subsets, including the increased ratios of neutrophil, NK cells, macrophage, and CD8 T cells, and the decreased ratios of B and CD4 T cells.

The applied antibody panels also included various other functional makers: antigen presentation and costimulation (major histocompatibility complex, MHC-II), cell activation and cell phenotype (CD44, IgM, Arg1), cell migration (CD62L), cell signaling transduction (CD38) and cytokine (IL-4, IL-6, IL-10, GM-CSF), tumor necrosis factor α(TNF-α), and interferon γ (IFN-γ) and immunosuppressive factor (CTLA-4, PD-1). A visualization technique, the heatmap, was then used to display the fold changes of functional markers in different cell subsets of the KO mice relative to those in the WT group (figure4E). The heatmap showed that while CD4+ T cells had higher expressions of Arg1, PD-1, and CD44, CD8+ T cells had higher expressions of CD44 and CD38 and the NK cells had a higher expression of CD38 (fold change>2). Meanwhile, fold changes of CTLA-4 in CD8+ T cells, CD62L in B cells, CD62L, CD44, MHCII, IL10, IL6 in macrophages, and CD44 and IL6 in neutrophil were decreased (fold change <0.5). However, we discovered that the magnitude of the difference in the expression level was quite small for most genes between the two groups. Nevertheless, these results showed that T cells and NK cells were activated in the Loxl1 KO mice, whereas B cells, macrophages, and neutrophils showed a reduction in migration and secretion acitivities in the KO mice group.

**Figure4.**
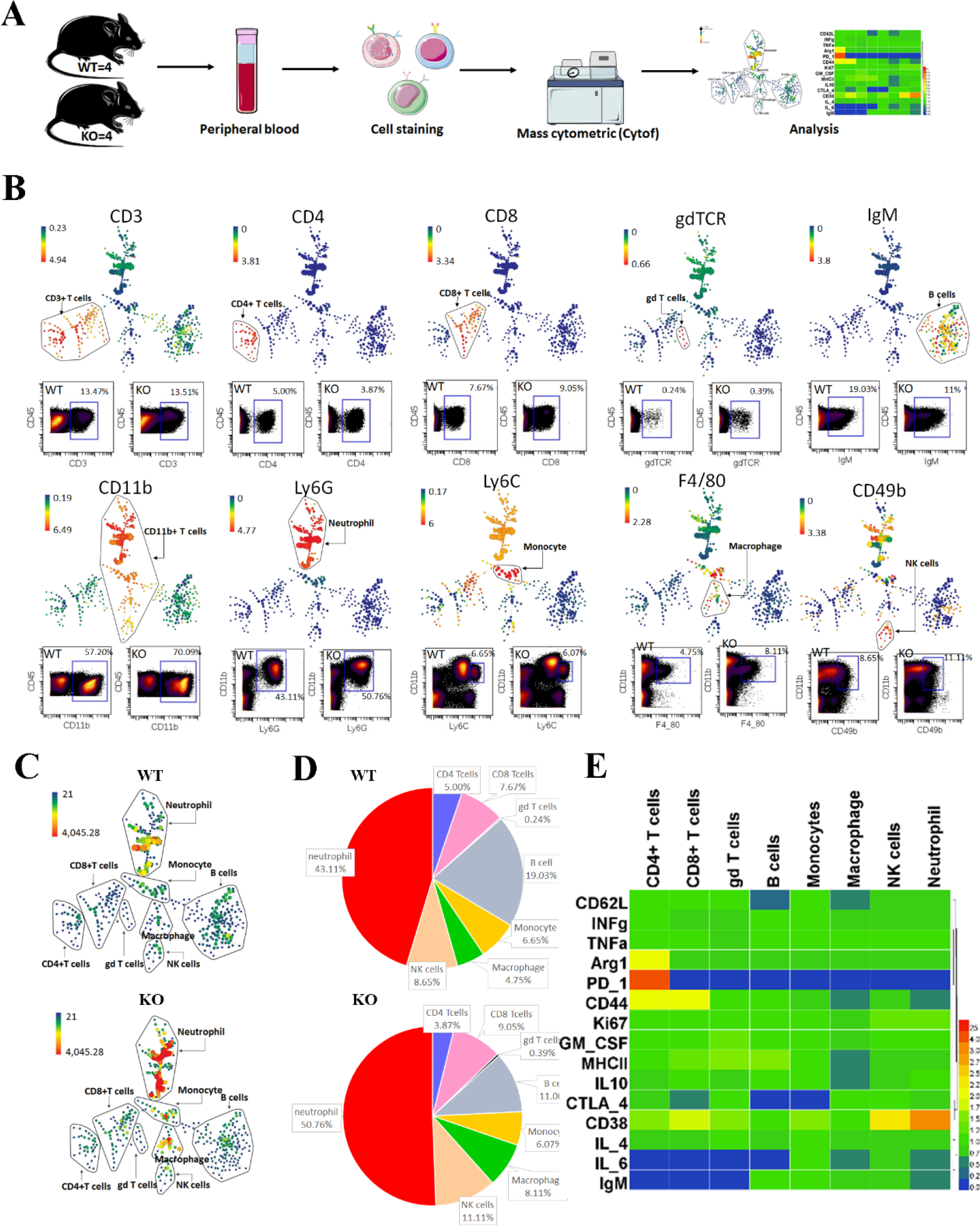
Systematic immune changes in KO mice. **A.** Scheme of the experimental procedure for characterization of immune cells populations in peripheral blood by mass cytometry. Peripheral blood extracted from 4 mice in each group were pooled as a single sample and stained with a cocktail of metal-tagged antibodys. Sample were analyzed using CyTOF machine and immune cell populations were identified using SPADE and in standard dot plots. **B.** SPADE analysis for characterizing the immune cell populations by using the canonical markers, including CD3, CD4, CD8, gdTCR, IgM, CD11b, Ly6G, Ly6C, F4/80 and CD49b. In SPADE, node size correlates with the number of cells, and the colored gradient corresponds to the arcsinh-transformed expression of the marker’s median. Dot plots demonstrated the marker labeling on CD45 positive cells and the proportion of each cell population in WT and KO groups were shown. **C.** SPADE diagram of the identified immune cell populations in WT and KO group. **D**. The relative abundance of identified immune cell populations in WT and KO group. **E.** Heatmap of the functional markers expression of immune cells. Values were foldchange of marker expression of immune cells in KO group compared to compartment in WT group.

**Supplement figure2.**
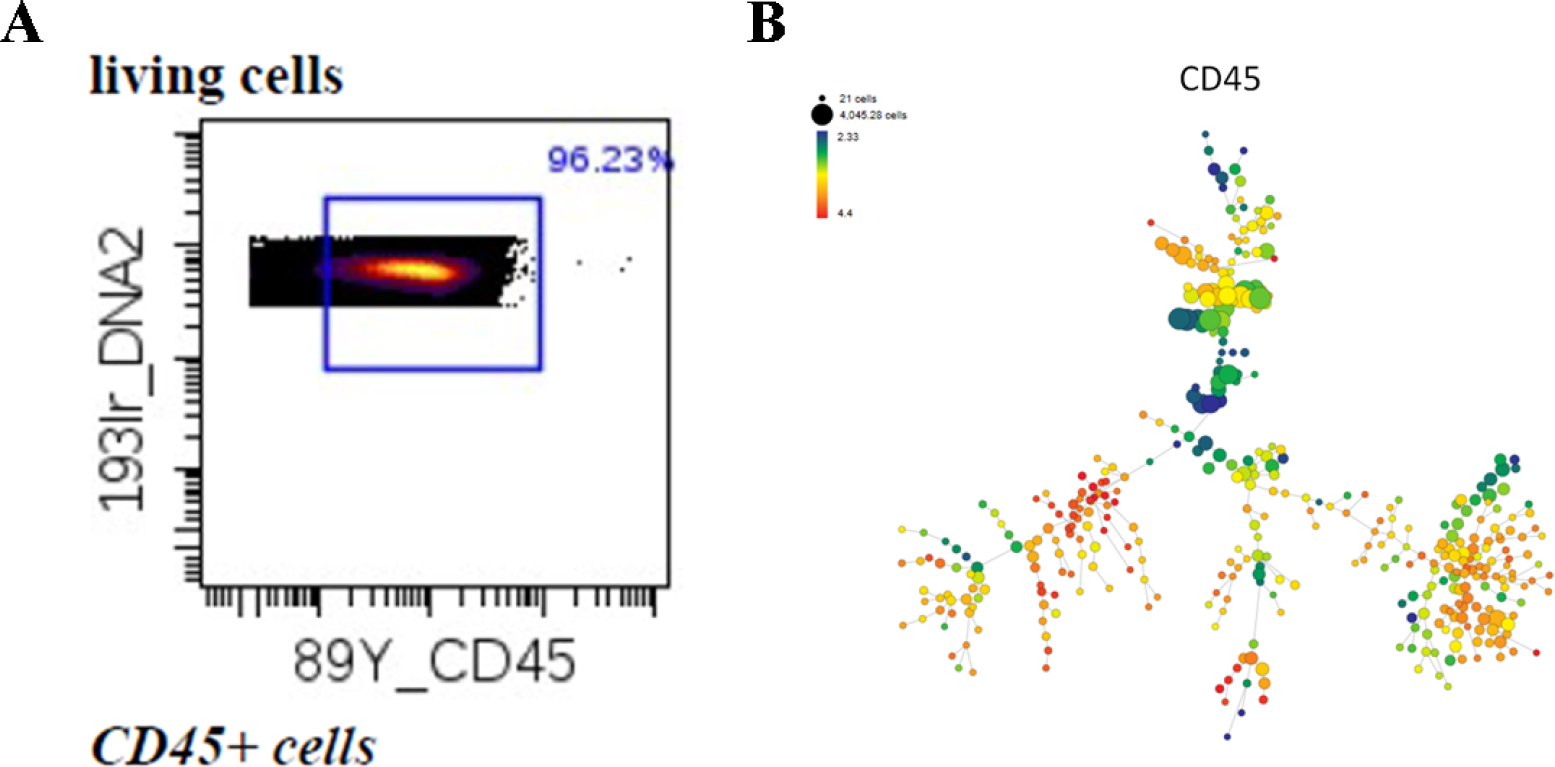
characterization of the immune cell populations by using the canonical markers CD45. **A** Dot plots demonstrated the marker labeling on live single cells and the proportion of CD45 positive cells in live single cells. **B** SPADE analysis for characterizing the overall expression of CD45 in immune cells.

### 5. Clinical relevance between *LOXL1* mutation and cancer

Due to the fact that *LOXL1* deficiency could induce changes in the immune system and hyper-proliferation in tissues, which are the two hallmarks often observed in tumors ^19^, we hypothesized that *LOXL1* mutation may be correlated with the progression of cancer. To test this hypothesis, we first collected data from The Exome Aggregation Consortium (ExAC, http://exac.broadinstitute.org/gene/ENSG00000129038) to identify *LOXL1* mutation in the common human population. According to the database, there were many forms of variations (including missense mutation, frameshift mutation, large fragment deletion, large fragment insertion, early termination, etc.) in the *LOXL1* gene in the population; missense mutation accounted for approximately 86% of the variations (Figure5A), leading to the disruption of polypeptide chain functions.

As the data set provided on ExAC spans 60,706,000 unrelated individuals, and can serve as a useful reference set of allele frequencies for severe disease studies, we next used the data set from ExAC as a control group representing the general population and collected gene mutation data set from TCGA and CanVar (https://canvar.icr.ac.uk/gene/ENSG00000129038) (two ontology database) as tumor patient groups. There were four common mutation loci of *LOXL1* in ExAC in the general population when compared with patients from TCGA and CanVar (figure5B). Additionally, the mutation frequency of the eight loci in tumor patients were significantly higher than the mutation frequency in the general population data set (figure5C, D), thus suggesting that higher frequency of *LOXL1* mutation may be related to tumor.

Next we compared the overall survival rate between the *LOXL1* non-mutant tumor patients and *LOXL1* mutant tumor patients from TCGA. In the pan-cancer patients, the overall survival rate of *LOXL1* mutant patients (n=15) were relatively lower when compared to the non-mutant patients (n=5780) (p=0.1438) (figure5E).

Results obtained from GO analysis showed that in liver hepatocellular carcinoma patients, cell cycle-related GO terms were significantly up-regulated in *LOXL1* mutant patients when compared with *LOXL1* non-mutant patients; similar outcomes were observed in GO terms related to mitotic cell cycle, mitotic cell cycle process, and cellular response to DNA damage (figure5F). In lung adenocarcinoma patients, immune activation-related GO terms including T cell proliferation, T cell migration, signal transduction, and cytokine production were significantly up-regulated in *LOXL1* mutant patients (figure5G). Therefore, when compared with *LOXL1* non-mutant patients, the mutant patients not only had a lower survival rate but also a higher transcriptomic level of genes related to biological processes such as cell proliferation and immune activation. These results were consistent with the phenotype identified in Loxl1 KO mice.

**Figure5.**
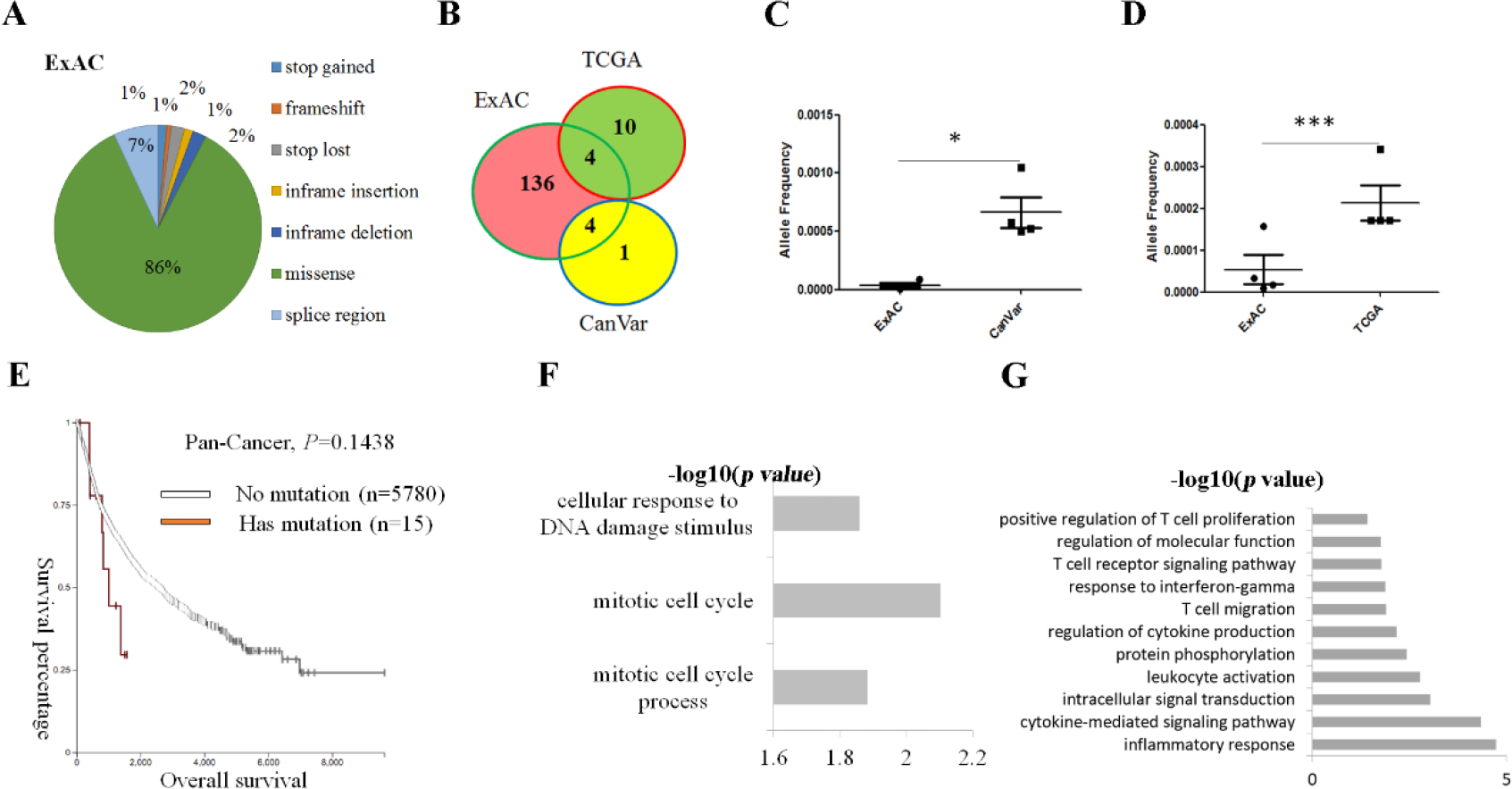
Clinical data about relationship between *LOXL1* mutation and cancer. **A.** Variation forms of *LOXL1* gene in population. **B.** Common mutation loci of *LOXL1* in ExAC in the general population when compared with cancer patients from TCGA and CanVar. **C, D.** The mutation frequency of the common mutation loci of *LOXL1* in ExAC in the general population when compared with cancer patients from CanVar (**C**) and TCGA (**D**) respectively. **E.** The overall survival rate of *LOXL1* mutant patients (n=15) when compared to the non-mutant patients (n=5780) in the pan-cancer patients. **F.** Biological process GO analysis of up-regulated genes in Loxl1 mutant patients with liver hepatocellular carcinoma using DAVID software. **G.** Biological process GO analysis of up-regulated genes in Loxl1 mutant patients with lung adenocarcinoma using DAVID software.

## Discussion

Our study exhibited that abnormal elastin activity induced by Loxl1 deficiency can result in a series of tissue-specific elastin-related modifications. By utilizing high-throughput sequencing combined with single cell CYTOF and histological evaluations, we demonstrated that Loxl1 deficiency in mice promoted local immune responses and hyperplasia in multiple tissues, as well as drastic changes in systemic immunity functions. Lastly, we also displayed the correlation between Loxl1 deficiency and tumorigenesis and poor survival rate in tumor patients.

ECM remodeling contributes to the progression and development of diseases ^4^ Our RNA-sequencing data showed that most of the significantly up-regulated DE genes (20/38) were related to extracellular matrix components (FBN1, COL6A3, COL1A2, COL1A1, LOXL1), suggesting that Loxl1 deficiency can induce elastin fragmentation and subsequent ECM remodeling. Results in our model also showed that elastin fragmentation and ECM remodeling resulted in the activation of major chemotactic activities, recruiting monocytes and macrophages to the local inflamed site in multiple organs, and initiating systemic immune response in the peripheral blood. Several publications illustrated that aberrant ECM expression and fragmentation of ECM can impact cell activation and survival, leading to inflammation and immune response activation ^20^. It was reviewed that the matrix fragments of collagen (the tripeptide N-acetyl Pro-Gly-Pro) are important for neutrophil chemo-attractants by mimic the chemotactic effects of CXCL8^21^. Other constituents of the ECM such as hyaluronan could induce the perpetuation of inflammatory responses through the activation of TLR2, or TLR4, or both ^22^; fibulin-5 can affect cutaneous inflammation through the regulation of active inflammatory level of NFkB ^23^. Thus, the enhancement of local immune infiltration of monocytes and macrophages in our Loxl1 KO mice may also be mediated through chemokine-and cytokine-induced immunity.

The immune system of our body always plays a crucial role in tissue homeostasis. For example, a large number of studies has proved that the activation of inflammatory responses is closely associated with tissue hyperplasia. Zhong et al. also illustrated that IL1 secreted by activated monocytes/macrophages can promote epithelial cell proliferation and increase epithelial layer thickness through paracrine action ^24^ Furthermore, in colitis, inflammatory mediators led to an increase in cell proliferation through the PI3K/Akt/β-Catenin pathway^25,26.^ The nuclear factor (NF) -κB family members and their inhibitors can maintain cell viability and proliferation through the IKK/IκB/NF-κB Pathway ^27^ Overall, these results help explain that the occurring hyperplasia in multiple tissues in the Loxl1 KO mice model may be mediated and followed by local immune infiltration and induction of systemic immunity, Our results also revealed a close relationship between Loxl1 mutation and tumor susceptibility, with a higher mutation frequency and lower overall survival rate in tumor patients with Loxl1 mutations. Inflamed environment and cell cycle phase were closely related to cancer progression and outcome ^28^ as the inflamed environment can disrupt the quiescence of tumor stem cells and initiate the tumorigenesis^29,30,^ which was consistent with our results that Loxl1 KO caused local and systematic immune responses as well as multiple tissue hyperplasia. Previous studies also illustrated that increased proliferation and self-renewal also play a key role in tumorigenesis; the increased DNA replication rate that occurs during proliferation makes it more susceptible to errors. Among all of the cancer-related mutations, 66.1% is due to random DNA replication errors ^31^. Therefore, further studies are required to identify whether Loxl1 knockout mice have higher random mutation rate.

In conclusion, we developed a strategy to combine single-cell mass cytometry and body-wide organ transcriptomic (BOT) analysis to systematically evaluate the effect of Loxl1 knockout on systemic immunity and tissue hyperplasia in mice. Thus, this research provided a powerful strategy to screen body-wide organ functions of a particular gene; the findings also illustrated the important biological roles of elastin on multiple organ cells and systemic immunity. These discoveries are both of important value for the understanding of ECM biology and pathology.

## Materials and Methods

### Animal model and body-wide organ collection

12-week-old Loxl1 knockout mice and C57BL/6 (both breed in National Resource Center for Mutant Mice, Model Animal Research Center of Nanjing University) were utilized in this study. Three animals respectively in KO mice group and WT mice group were humanely sacrificed and the 17 tissues were collected for subsequent experiments. Four animals in each group were humanely sacrificed and the peripheral blood were pooled into one sample for single cell mass cytometry. All procedures were performed with approved protocols, in accordance with guidelines of the animal experimental center of Zhejiang University, People’s Republic of China.

### Histological examination

Specimens were immediately fixed in 4% neutral buffered paraformaldehyde, dehydrated through an alcohol gradient, cleared, and embedded in paraffin blocks. Histological sections (7 μm) were prepared using a microtome, and subsequent lyde-paraffinized with xylene, hydrated using decreasing concentrations of ethanol and then subjected to hematoxylin and eosin (H&E) staining and Weigarts’ staining. Then the sections were mounted and observed under microscopy.

### RNA-seq

RNA-seq was modified from a previous method ^32^ Briefly, RNA was extracted from samples by Trizol reagent (TAKARA), reverse transcription was conducted by SuperScript II reverse transcriptase (Invitrogen), double strand cDNA was conducted using NEBNext mRNA second strand synthesis kit (NEB), double strand DNA was cleaned with AMPure XP beads (Beckman Coulter), sequencing library was constructed with Nextera XT kit(Illumina) and sequenced on Illumina X-Ten platform. RNA-seq reads data were mapped to reference genome using TopHat and Cufflinks ^33^. Expression was calculated with counts per million (CPM).

### Data analysis for RNA-seq

Differential expression gene was calculated by DESeq2, differential expressed genes were selected with *P* value<0.05 ^34^ Gene ontology enrichment analysis were performed with DAVID informatics resources (https://david.ncifcrf.gov/) ^35^.

### Immunofluorecence staining

The tissues were fixed in 4% (w/v) paraformaldehyde, and then dehydrated in an ethanol gradient, prior to embedment in paraffin and sectioning at 7μm thickness. Immunostaining were carried out as follows: The 7μm paraffin sections were rehydrated, antigen retrieved, rinsed three times with PBS, and treated with blocking solution (1% BSA) for 30 min, prior to incubation with primary antibodies at 4 °C overnight. The primary antibodies: rabbit anti-mouse antibodies against KI67 (Abcam, ab16667) and CD45 (BD Biosciences, 555483), rat anti-mouse CD19 (Biolegend, 115525) and F4/80 (Biolegend, 123121) were used to detect the cell proliferation and immune cell infiltration. Secondary antibody: goat anti-rabbit Alexa Fluor 488 (Invitrogen, A11008), goat anti-rabbit Alexa Fluor 546 (Invitrogen, A21430-f), goat anti-rat CY3 (Beyotime Institute of Biotechnology, A0507), and DAPI (Beyotime Institute of Biotechnology, China) were used to visualize the respective primary antibodies and the cell nuclei. All procedures were carried out according to the manufacturer’s instructions.

### Single cell mass cytometry

We pooled the blood from four mice for each group to obtain sufficient cells for a reliable mass cytometry. After lysis of the erythrocyte by ACK lysis buffer, samples were washed by FACS buffer and kept at 4 °C or on ice. The pooled cells were then stained with cisplatin for live-dead cell distinction. Following blocked at room temperature for 20 min, cells were stained with a mixture of metal-tagged antibodies targeting the surface antigens for 30 min at room temperature (a complete list of antibodies is provided in Supplementary Table 1). Following washed with FACS buffer and fixed for 20 min at RT, cells were washed with Perm buffer and stained with a mixture of intracellular antibodies (Supplementary Table 1) for 30 min at RT. Then the cells were fixed and stained with DNA intercalator overnight at 4°C. After washed by perm buffer, cells were incubated with barcodes at 4 °C for 30min. Following multiple washes with FACS buffer and ultrapure H_2_O, cells were analyzed on a CyTOF machine. The raw data acquired was uploaded to a Cytobank web server (Cytobank Inc.) for further data processing and for gating out of dead cells and normalization beads.

**Supplement table 1:**
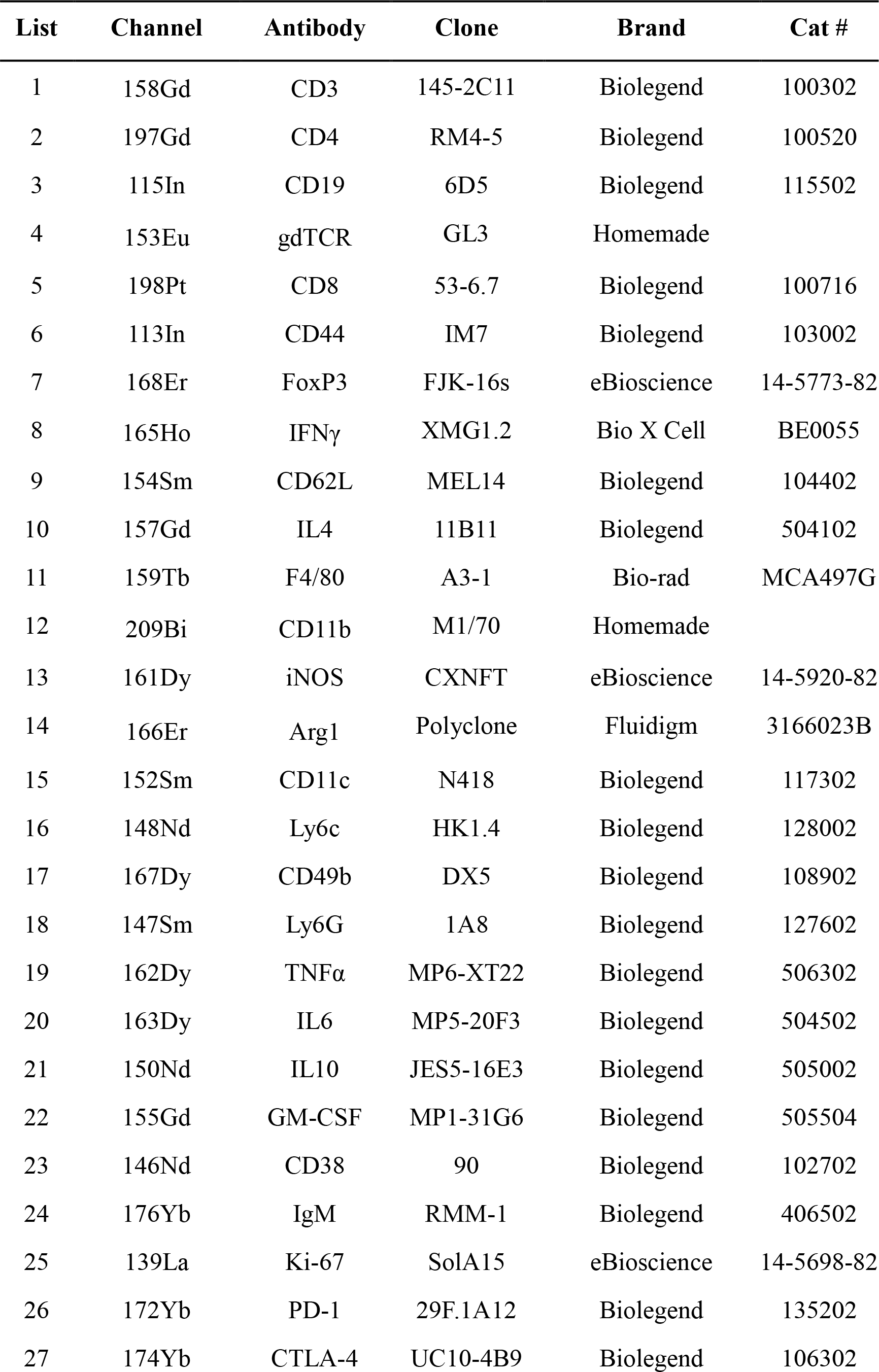
Antibodies used in CyTOF.

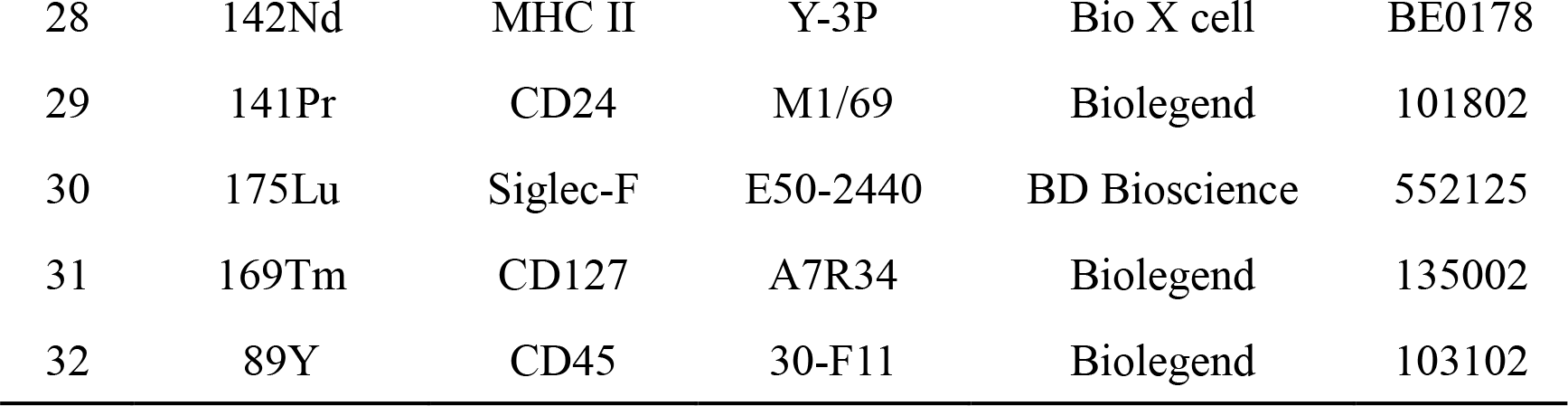

### Mass Cytometry Data analysis

Data analysis was performed using SPADE ^36^ algorithms and scatter diagram in Cytobank (www.cytobank.org). SPADE analyses in Cytobank were performed with the target number of nodes set to 200. In SPADE algorithms, the color gradient represents the median expression level of the chosen marker. The following markers were chosen for immune-related clustering: CD45, CD3, CD4, CD8, gdTCR, IgM, CD11b, Ly6G, Ly6C, F4/80 and CD49b. Cell populations were then identified based on marker expression distribution in the figure. The percentage of each immune cell population was calculated for each defined population out of the total CD45 positive cells. Intensity levels of markers were calculated on transformed median intensity values in each defined cell population. Heatmaps were drawn using HemI 1.0.3.3 Heatmap Illustrator ^37^.

### Statistical analysis

Two-tailed t-test was used to detect differences of histological results between the KO mice and WT mice groups. The statistical significance set at p <0.05.

## Acknowledgements

This work was supported by the National High Technology Research and Development Program of China (2016YFB0700804), the National Natural Science Foundation of China (CN) (81270682, 81300454), the Key Scientific and Technological Innovation Team of Zhejiang Province (2013TD11).

## Author contributions

B.B.W.: data, data analysis and interpretation, manuscript writing. Y.L.: data, sample processing for histology and immunofluorescence, manuscript writing; C.R.A.: animal experiments, sample processing; D.M.J., L.G., Y.S.L., Y.X.L,: sample collection; J.L.: figure reorganize; H.W.O.Y.: conception and design, manuscript writing; X.H.Z.: conception and design, manuscript writing.

## References

1 Hynes, R. O. The extracellular matrix: not just pretty fibrils. Science 326, 1216–1219, doi:10.1126/science.1176009 (2009).

2 Calve, S., Odelberg, S. J. & Simon, H. G. A transitional extracellular matrix instructs cell behavior during muscle regeneration. Developmental Biology 344, 259–271, doi:10.1016/j.ydbio.2010.05.007 (2010).

3 Bassat, E. et al. The extracellular matrix protein agrin promotes heart regeneration in mice. Nature 547, 179−+, doi:10.1038/nature22978 (2017).

4 Bonnans, C., Chou, J. & Werb, Z. Remodelling the extracellular matrix in development and disease. Nat Rev Mol Cell Biol 15, 786–801, doi:10.1038/nrm3904 (2014).

5 Guilak, F. et al. Control of Stem Cell Fate by Physical Interactions with the Extracellular Matrix. Cell Stem Cell 5, 17–26, doi:10.1016/j.stem.2009.06.016 (2009).

6 Mammoto, A. et al. Control of lung vascular permeability and endotoxin-induced pulmonary oedema by changes in extracellular matrix mechanics. Nat Commun 4, doi:ARTN 175910.1038/ncomms2774 (2013).

7 Rozario, T. & DeSimone, D. W. The extracellular matrix in development and morphogenesis: a dynamic view. Developmental biology 341, 126–140, doi:10.1016/j.ydbio.2009.10.026 (2010).

8 Nakasaki, M. et al. The matrix protein Fibulin-5 is at the interface of tissue stiffness and inflammation in fibrosis. Nature communications 6, 8574, doi:10.1038/ncomms9574 (2015).

9 Chen, H., Zheng, X. & Zheng, Y. Age-associated loss of lamin-B leads to systemic inflammation and gut hyperplasia. Cell 159, 829–843, doi:10.1016/j.cell.2014.10.028 (2014).

10 De Arcangelis, A. et al. Hemidesmosome integrity protects the colon against colitis and colorectal cancer. Gut 66, 1748–1760, doi:10.1136/gutjnl-2015-310847 (2017).

11 Hynes, R. O. & Naba, A. Overview of the matrisome--an inventory of extracellular matrix constituents and functions. Cold Spring Harbor perspectives in biology 4, a004903, doi:10.1101/cshperspect.a004903 (2012).

12 Tassabehji, M. et al. An elastin gene mutation producing abnormal tropoelastin and abnormal elastic fibres in a patient with autosomal dominant cutis laxa. Human molecular genetics7, 1021–1028 (1998).

13 Morris, C. A. & Mervis, C. B. Williams syndrome and related disorders. Annual review of genomics and human genetics 1, 461–484, doi:10.1146/annurev.genom.1.1.461 (2000).

14 Liu, X. et al. Elastic fiber homeostasis requires lysyl oxidase-like 1 protein. Nature genetics 36, 178–182, doi:10.1038/ng1297 (2004).

15 Kagan, H. M. & Li, W. Lysyl oxidase: properties, specificity, and biological roles inside and outside of the cell. Journal of cellular biochemistry 88, 660–672, doi:10.1002/jcb.10413 (2003).

16 Molnar, J. et al. Structural and functional diversity of lysyl oxidase and the LOX-like proteins. Biochimica et biophysica acta 1647, 220–224 (2003).

17 Proserpio, V. & Lonnberg, T. Single-cell technologies are revolutionizing the approach to rare cells. Immunol Cell Biol 94, 225–229, doi:10.1038/icb.2015.106 (2016).

18 Drewes, P. G. et al. Pelvic organ prolapse in fibulin-5 knockout mice: pregnancy-induced changes in elastic fiber homeostasis in mouse vagina. The American journal of pathology 170, 578–589, doi:10.2353/ajpath.2007.060662 (2007).

19 Hanahan, D. & Weinberg, R. A. Hallmarks of cancer: the next generation. Cell 144, 646–674, doi:10.1016/j.cell.2011.02.013 (2011).

20 Sorokin, L. The impact of the extracellular matrix on inflammation. Nat Rev Immunol 10, 712–723, doi:10.1038/nri2852 (2010).

21 Weathington, N. M. et al. A novel peptide CXCR ligand derived from extracellular matrix degradation during airway inflammation. Nat Med 12, 317–323, doi:10.1038/nm1361 (2006).

22 Brusselle, G. G., Joos, G. F. & Bracke, K. R. Chronic Obstructive Pulmonary Disease 1 New insights into the immunology of chronic obstructive pulmonary disease. Lancet 378, 1015–1026, doi: Doi 10.1016/S0140-6736(11)60988-4 (2011).

23 Nakasaki, M. et al. The matrix protein Fibulin-5 is at the interface of tissue stiffness and inflammation in fibrosis. Nature communications 6, doi:ARTN 857410.1038/ncomms9574 (2015).

24 Zhong, F. L. et al. Germline NLRP1 Mutations Cause Skin Inflammatory and Cancer Susceptibility Syndromes via Inflammasome Activation. Cell 167, 187–202 e117, doi:10.1016/j.cell.2016.09.001 (2016).

25 Kaler, P., Godasi, B. N., Augenlicht, L. & Klampfer, L. The NF-kappaB/AKT-dependent Induction of Wnt Signaling in Colon Cancer Cells by Macrophages and IL-1beta. Cancer Microenviron 2, 69–80, doi:10.1007/s12307-009-0030-y (2009).

26 Lee, G. et al. Phosphoinositide 3-Kinase Signaling Mediates beta-Catenin Activation in Intestinal Epithelial Stem and Progenitor Cells in Colitis. Gastroenterology 139, 869–U237, doi:10.1053/j.gastro.2010.05.037 (2010).

27 Ghosh, S., May, M. J. & Kopp, E. B. NF-kappa B and Rel proteins: evolutionarily conserved mediators of immune responses. Annu Rev Immunol 16, 225–260, doi:10.1146/annurev.immunol.16.1.225 (1998).

28 Gentles, A. J. et al. The prognostic landscape of genes and infiltrating immune cells across human cancers. Nat Med 21, 938–945, doi:10.1038/nm.3909 (2015).

29 Moon, H. et al. Melanocyte Stem Cell Activation and Translocation Initiate Cutaneous Melanoma in Response to UV Exposure. Cell stem cell 21, 665–678 e666, doi:10.1016/j.stem.2017.09.001 (2017).

30 Westphalen, C. B. et al. Dclk1 Defines Quiescent Pancreatic Progenitors that Promote Injury-Induced Regeneration and Tumorigenesis. Cel stem cell 18, 441–455, doi:10.1016/j.stem.2016.03.016 (2016).

31 Tomasetti, C., Li, L. & Vogelstein, B. Stem cell divisions, somatic mutations, cancer etiology, and cancer prevention. Science 355, 1330–1334, doi:10.1126/science.aaf9011 (2017).

32 Picelli, S. et al. Smart-seq2 for sensitive full-length transcriptome profiling in single cells. Nat Methods 10, 1096–1098, doi:10.1038/nmeth.2639 (2013).

33 Trapnell, C. et al. Differential gene and transcript expression analysis of RNA-seq experiments with TopHat and Cufflinks. Nat Protoc 7, 562–578, doi:10.1038/nprot.2012.016 (2012).

34 Love, M. I., Huber, W. & Anders, S. Moderated estimation of fold change and dispersion for RNA-seq data with DESeq2. Genome Biol 15, 550, doi:10.1186/s13059-014-0550-8 (2014).

35 Huang, D. W., Sherman, B. T. & Lempicki, R. A. Systematic and integrative analysis of large gene lists using DAVID bioinformatics resources. Nature Protocols 4, 44–57 (2009).

36 Qiu, P. et al. Extracting a cellular hierarchy from high-dimensional cytometry data with SPADE. Nat Biotechnol 29, 886–U181 (2011).

37 Deng, W. K., Wang, Y. B., Liu, Z. X., Cheng, H. & Xue, Y. HemI: A Toolkit for Illustrating Heatmaps. Plos One 9 (2014).

